# A novel proneural function of Asense, integrated with of L’sc and Notch, promotes the Neuroepithelial to Neuroblast transition

**DOI:** 10.1101/2021.04.15.440037

**Authors:** Mercedes Martin, Mirja N. Shaikh, Francisco Gutierrez-Avino, Francisco J. Tejedor

## Abstract

In the developing *Drosophila* optic lobe, neuroepithelial (NE) cells are transformed progressively into neurogenic progenitors called neuroblasts (NBs). The progenitors undergoing this transition are identified by the expression of the Acheate Scute Complex (AS-C) factor Lethal of Scute (L’sc).

Here we found that Asense (Ase), another AS-C factor, presents a peak of expression in the cells neighboring those transition L’sc expressing cells. This peak of Ase identifies a new transition step and it is necessary and sufficient to promote the NE to NB transition. Thus, our data provide the first direct evidence for a proneural role of Ase in CNS neurogenesis. Furthermore, we found that the peak of Ase is induced in a non-cell autonomous manner by L’sc through the activation of Notch signaling in the adjacent cells. This suggests that the two classic proneural activities, promoting neurogenesis and Notch signaling, have been split between Ase and L’sc. Thus, our data fit with a model in which the key proneural role of Ase is integrated with Notch and L’sc activities, facilitating the progressive transformation of NE cells into NBs.

**SUMMARY STATEMENT:** The switch of progenitor cells towards neuron production is crucial for proper brain development. The transcription factor Asense promotes this transition through a mechanism integrated with timing and neurogenic signals.

## INTRODUCTION

The correct formation of a functional nervous system depends on the dynamic coordination of NPs proliferation and differentiation. During the development of a complex CNS, NPs progress sequentially through distinct stages, initially dividing symmetrically to expand their population and later, dividing asymmetrically to produce different neuronal populations in a stepwise manner (reviewed by Caviness et al., 1995; Götz & Huttner, 2005; Kriegstein et al., 2006; Perry et al., 2017). This transition from proliferating to neurogenic NPs is a key step that must be tightly regulated during neurogenesis (reviewed by Huttner & Brand, 1997; Huttner & Kosodo, 2005; Kriegstein et al., 2006). Hence, it is important to unravel the full set of genes and molecular mechanisms that control this process.

The larval Optic Lobe (OL) primordia of *Drosophila melanogaster (Dm)* is an experimental model very well suited to analyze the genetic and molecular basis of this transition, particularly given the well-defined spatio-temporal organization of these cellular processes and the availability of genetic tools (Brand & Livesey, 2011; Ceron et al., 2001; Egger et al., 2007, 2011; Homem & Knoblich, 2012; Perry et al., 2017; Sousa-Nunes et al., 2010; Suzuki & Sato, 2014). The *Dm* OL originates from an epithelial vesicle that invaginates from the head epidermis and becomes attached to the brain (Green et al., 1993; Namba & Minden, 1999). This small cluster of NE cells reactivates to increase in number during 1^st^ and 2^nd^ instar larvae, and it becomes segregated into two primordia by the end of this period: the Outer Proliferation Center (OPC) and the Inner Proliferation Center (IPC). Of these, the OPC gives rise to the precursor cells of the lamina and the outer medulla, while the IPC generates those of the lobula complex and the inner medulla (Hofbauer & Campos-Ortega, 1990; Meinertzhagen & Hanson, 1993; White & Kankel, 1978).

These anlagen initially grow as a result of the symmetric division of NE cells. Yet during the 3^rd^ larval instar, OPC NE cells progressively differentiate into medulla NBs and lamina precursor cells at the medial and lateral edges, respectively. The medulla NBs then begin to divide asymmetrically in a self-renewing fashion producing a new NB and a ganglion mother cell (GMC) that in turn divides once into two medulla neuronal precursors called ganglion cells (GCs) (Ceron et al., 2001). These sequential processes are reminiscent of those that happen in the developing mammalian cerebral cortex and neural tube (Brand & Livesey, 2011; Doe, 2008; Homem et al., 2015; Suzuki & Sato, 2017). In these mammalian tissues, the NE cells in the ventricular zone (VZ) initially divide symmetrically to expand the pool of progenitors and then, at the onset of neurogenesis, they progressively lose their NE properties and are transformed into radial glial cells that divide asymmetrically to generate intermediate progenitors witch move to the subventricular zone (SVZ) where they terminally divide into two neurons (reviewed by Kriegstein et al., 2006).

The NE to NB transition in the OPC progresses in a spatially and temporally ordered manner in a medial to lateral direction (Egger et al., 2007). This stepwise transition is crucial to progressively generate the different populations of medulla neurons (Li et al., 2013; Sato et al., 2013) and it requires the coordinated action of several signaling pathways, mainly JAK/STAT, EGFR, Fat-Hippo and Notch (reviewed by Apitz & Salecker, 2014; Egger et al., 2011; Sato et al., 2013). Thus, JAK/STAT signaling is activated in the lateral NE cells of the OPC, initially promoting NE expansion and negatively regulating the progression of the NE-NB transition (Ngo et al., 2010; Wang et al., 2011a; Yasugi et al., 2008). EGFR signaling is required for the proliferation of NE cells and to induce the expression of the L’sc proneural factor (Yasugi et al., 2010) that precedes NB formation (Yasugi et al., 2008). L’sc promotes cell-autonomously the expression of Delta (Dl), which in turn activates the Notch pathway in adjacent cells on both the medial (NB) and lateral (NE) sides. While this oscillation of Notch signaling seems to be required for the transition to NBs, the activation of Notch in the NEs represses *l’sc* expression and prevents a premature switch to NBs (Egger et al., 2010; Ngo et al., 2010; Reddy et al., 2010; Wang et al., 2011b; Weng et al., 2012; Yasugi et al., 2010). In this way, the coordination of these signals produces a wave of *l’sc* expression (the proneural wave: Yasugi et al., 2008) that sweeps across the NE in a medial to lateral direction (Sato et al., 2016; Yasugi et al., 2010).

Although it has been reported that *l’sc* expression is sufficient to induce the NB fate (Egger et al., 2010; Yasugi et al., 2008), its loss of function does not prevent the NE-NB transition but rather, it simply alters the timing of NB differentiation (Yasugi et al., 2008). Moreover, it remains unclear how L’sc can induce the NB fate. L’sc belongs to the Achaete Scute Complex (AS-C) family of bHLH transcription factors (TFs), the products of the *achaete (ac), scute (sc), l′sc,* and *asense (ase)* genes (Campuzano et al., 1985; García-Bellido, 1979; Ghysen & Dambly-Chaudière, 1988; Romani et al., 1989). Classic studies of these genes showed they are required for the commitment of epithelial cells to the NP lineage in the peripheral nervous system (PNS) and CNS of *Dm* (reviewed by Campos-Ortega, 1993; García-Bellido & de Celis, 2009; Jan & Jan, 1993).

There is some redundancy in the roles of the AS-C TFs in the NE-NB transition (Yasugi et al., 2008). In addition to L’sc, other AS-C proteins are expressed in the OL, with *sc* expressed in NE cells and NBs (Egger et al., 2007), and *ase* in NBs and their GC progeny (Brand et al., 1993; González et al., 1989; Shaikh et al., 2016; Yasugi et al., 2008). Remarkably, *ase*-deficient flies develop a defective adult OL (González et al., 1989), although it has been reported that the deletion of *ase* alone does not overtly affect NB formation. Nevertheless, the delay in NB formation associated to the loss of *l′sc* is enhanced by the additional deletion of *ase* (Yasugi et al., 2008).

Ase plays an important role in regulating proliferation in the OL (Shaikh et al., 2016; Wallace et al., 2000), particularly in terms of GC cell cycle exit and terminal differentiation (Shaikh et al., 2016). However, its role in NB formation remains unclear and no proneural role has so far been reported in the CNS. Notably, DamID analysis and expression profiling of *ase* mutant embryos predict a dual role for Ase. On the one hand, Ase could activate the expression of self-renewal genes and repress the expression of differentiation genes in NBs, while on the other hand, it could promote the neuronal differentiation of the NB progeny (Southall & Brand, 2009). Interestingly, Ase plays a proneural role in the generation of bristles at the anterior wing margin (Domínguez & Campuzano, 1993). Moreover, Ascl1/Mash1, its closest vertebrate orthologue, plays an important proneural role in CNS neurogenesis (reviewed by Bertrand et al., 2002; Castro et al., 2011; Guillemot & Hassan, 2017; Vasconcelos & Castro, 2014).

In the light of these premises, we decided to assess a possible role of Ase in the NE to NB transition. Accordingly, we studied the expression of Ase during this transition in detail, and the possible alterations that its gain and loss of function (GoF/LoF) cause in those cellular processes in comparison with equivalent genetic conditions of L’sc. Furthermore, we analyzed how the expression of Ase is regulated, as well as the integration of Ase functions with L’sc and Notch signaling in this context. Additionally, we discuss the possible proneural functions of Ase in the Dm OL compared with those of its vertebrate orthologue Ascl1/Mash1.

## RESULTS

### *ase* expression peaks at the NE-NB transition

Ase has been commonly used as a molecular marker for central brain (CB) type I and OPC NBs in the larval brain (Egger et al., 2007; Orihara-Ono et al., 2011; Yasugi et al., 2008). Nevertheless, we previously found that *ase* expression is transiently upregulated in newborn GCs in which it is expressed more strongly than in the dividing NBs (Shaikh et al., 2016; Shaikh & Tejedor, 2018). Moreover, in those studies we also noted differences in *ase* expression level among the OPC NBs.

To more precisely define the expression of *ase* during the NE-NB transition, we assessed Ase immunostaining in *wt* brains in relation to the distribution of markers for NE cells (DE-Cadherin, Cad), transition cells (L’sc), and NBs (Deadpan, Dpn; Miranda, Mira) (Fig. 1B-G, Fig. 2A,C). As described previously (Brand et al., 1993; Wallace et al., 2000), Ase was found in all NBs but not in NEs or transition (L’sc^+^) cells. However, it was notable that the cells neighboring the L’sc^+^ transition cells were more strongly stained for Ase than the subsequent NBs (Fig. 1B-E). Moreover, these cells strongly stained for Ase expressed NB markers weakly or not at all (Fig. 1C,F and Fig. 2C), suggesting that these cells might still be in a transition phase. Henceforth, we will refer to these high-level Ase-expressing cells (i.e. strong initial upregulation of *ase*) as the “peak of Ase” cells.

**Figure 1.**
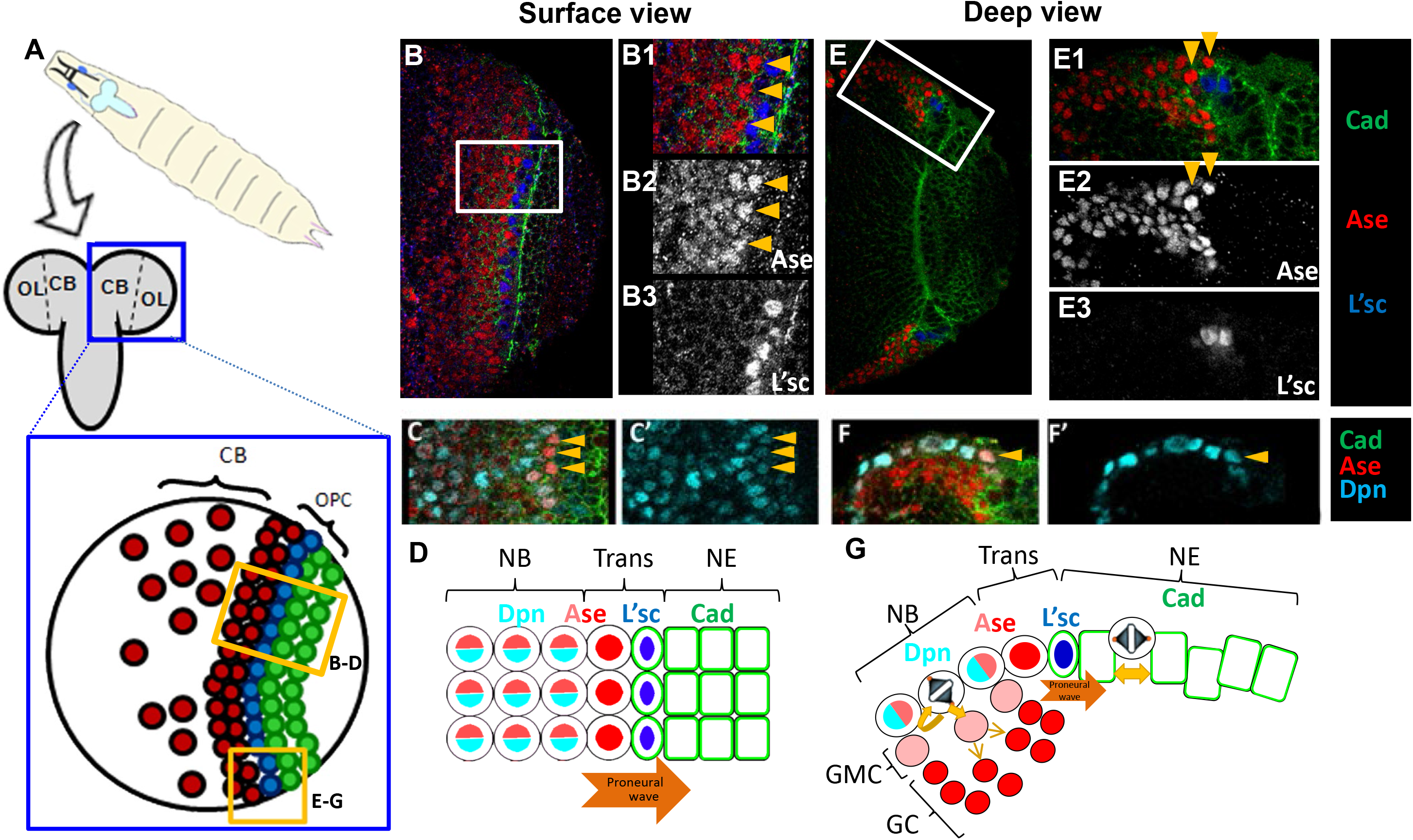
Expression of Ase in the OPC. **A.** Scheme representing the ventral view of the larval *Dm* brain indicating the location of the CB, OL and OPC regions, as well as the position of the NE cells (green), NE-NB transition cells (blue) and NBs (red) within the OPC. **B, C, E, F**. Confocal images showing the expression of Ase, Cad, Dpn and L’sc in the surface and deep layers, and high magnification views of the boxed areas as indicated. Note that the strongest Ase immunostained cells are neighbors of the L’sc^+^ cells and they have little Dpn (arrowheads). **D, G**. Scheme summarizing the data depicted in the preceding panels.

**Figure 2.**
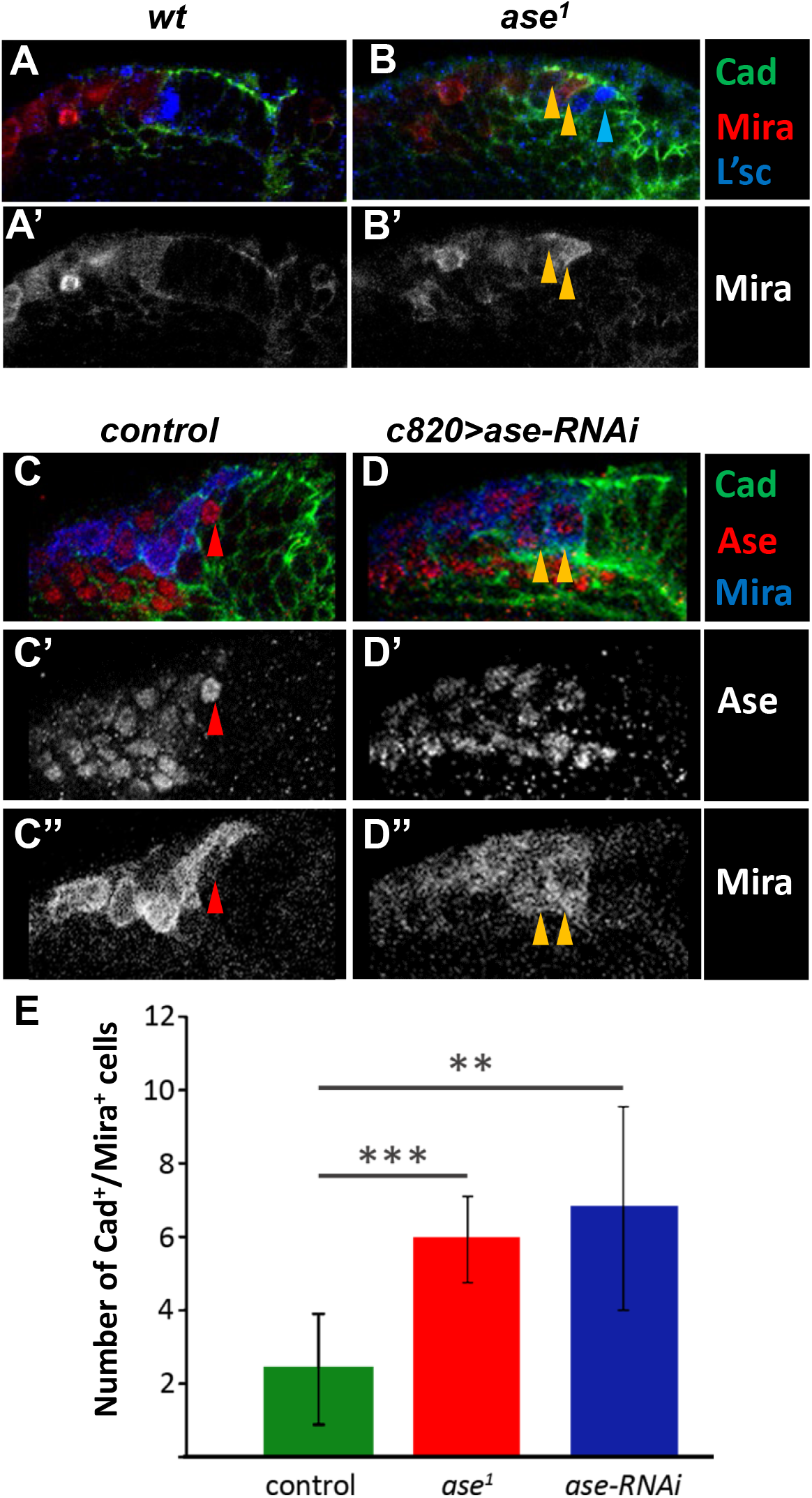
Alterations to the NE-NB transition by *ase LoF*. **A-D**: High magnification confocal images of deep OPC layers of *wt*, *ase*^*1*^, *c820 Gal4 (control)* and *c820 Gal4; UAS ase-RNAi* flies showing the staining for Ase, Cad, Mira and L’sc as indicated. **A, B**: Note that in the *ase*^*1*^ larvae there are cells co-expressing Cad and Mira (yellow arrowheads) whereas they are segregated in control samples. In addition, the *ase*^*1*^ sample has L’sc stained cells in the middle of the NE (blue arrowhead) instead of at its edge, as in the *wt* larvae. *C, D*. The *c820 >ase-RNAi* sample lacks cells with peak of *ase* expression relative to the controls (red arrowheads) and they have several cells co-expressing Mira and Cad (yellow arrowheads). **E**. Quantification of the Mira/Cad co-immunostained cells in control (n=24), *ase*^*1*^ (n=32) and *c820 > ase-RNAi* (n=10) OLs. Statistical significance was assessed with the student ***t***-test: control *vs*. *ase*^*1*^ = 5.5 × 10^-^7 and control *vs*. *ase-RNAi* = 3.5 = 10^-3^.

### Ase is necessary for proper NE-NB transition

We wondered if this oscillation of *ase* expression might be implicated in the NE to NB transition. To investigate this possibility, we analyzed the expression of differential markers of the NE to NB transition under *ase* GoF and LoF conditions. We first studied *ase*^*1*^, a 17 kb deletion of the whole gene that does not affect any other component of the AS-C (González et al., 1989) and we found that Mira (NB marker) and Cad (NE marker) were co-expressed in many cells during the NE-NB transition compared to *wt* brains (Fig. 2A,B and E). Furthermore, although L’sc was restricted to only a few cells in this mutant, as in the *wt* (Fig. 2A,B), many of these cells were located in the middle of the NE rather than at its edge (Fig. 2B). Hence, Ase appears to be required for the correct NE-NB transition.

We next asked whether the peak of Ase expression is specifically needed for the correct NE-NB transition, taking advantage of the *c820-Gal4* line (Manseau et al., 1997) that drives expression in the most medial NE and transition zone cells (Suppl. Fig. 1A). Thus, to eliminate the peak of Ase, *ase-RNAi* was expressed with this Gal4 line and we indeed observed the absence of high-level Ase-expressing cells near the NE. Moreover, in this condition, all Ase^+^ cells at the surface of the OPC exhibited weak Ase labeling and co-expressed Mira (Fig. 2C,D). Remarkably, this *ase-RNAi* condition caused the same phenotype as *ase*^*1*^, with a significant increase in the number of cells co-expressing NE and NB markers (Fig. 2B,D and E). Hence, we concluded that the peak of Ase is crucial for the correct NE-NB transition.

### Ase promotes the NE-NB transition

To further analyze the implication of Ase in the NE-NB transition, we misexpressed *ase* in the NE using the *c855a-Gal4* driver that directs expression to the OPC and IPC NE cells (Hrdlicka et al., 2002; Suppl. Fig. 1B). First, we drove *ase* expression from the beginning of the 3^rd^ instar (see Materials and Methods), a period when the NE to NB transition has barely started. This induced a decrease in the size of the OL and remarkably, the almost complete elimination of OPC NE cells at the expense of NBs, while GCs were located normally inside the OPC (Fig. 3A,B). These data strongly suggest that the misexpression of *ase* in NE cells induces the premature formation of NBs, thereby reducing the pool of NE cells and consequently, the size of the OL.

**Figure 3.**
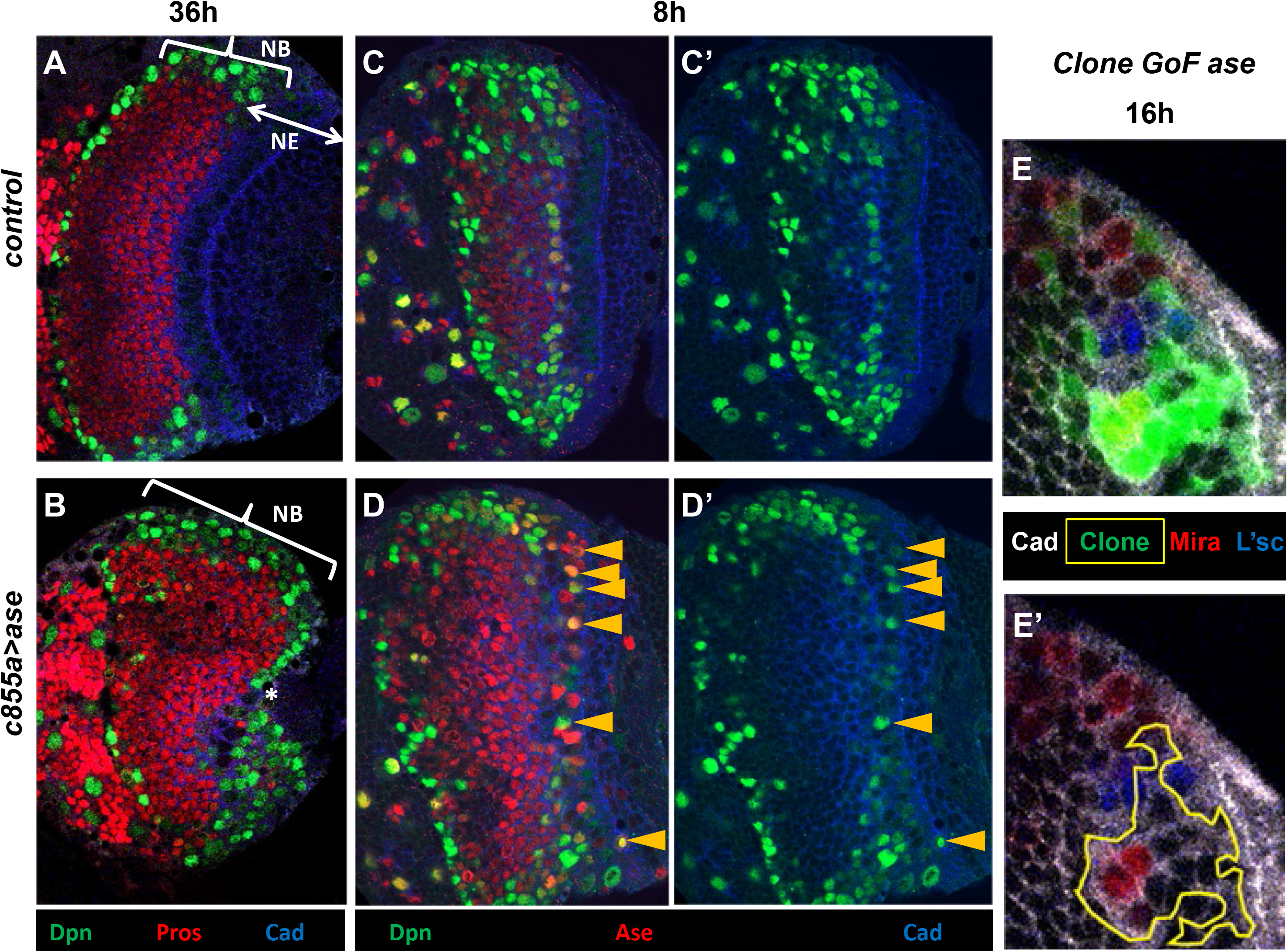
Misexpression of Ase in the NE generates NBs. **A, B**. Confocal images close to the surface of the OPC of control (*c855a Gal4*) and *c855a Gal4/UAS ase* larval brains after a 36h induction. Note that the OL size is reduced in the *c855a> ase* specimens relative to the controls and that there are no NE cells (Cad^+^ cells). Moreover, the most lateral region (NE in the control sample) is occupied by NBs (Dpn^+^) in the *c855a>ase* sample while in both specimens the interior area of the OPC is full of GCs (Pros^+^). **C, D**. Similar experiment after an 8h induction. Note the numerous intermingled Dpn^+^/Ase^+^ cells in the NE (yellow arrowheads) of the *c855a>ase* larvae relative to the controls. **E, E**’. Ase misexpressing clones originated in the NE (notice the apical Cad labeling) contain several Mira^+^ cells inside the OPC but do not exhibit L’sc immunostaining.

To assess if Ase could directly promote the transformation of NE cells into NBs, A*se* was misexpressed in larvae for a short period of time (8h) at the end of 3^rd^ instar using the same *c855a-Gal4* line. Interestingly, this condition produced ectopic cells expressing NB markers (Dpn and Mira) within the NE, where Ase was ectopically induced, and at its edge (Fig. 3C,D and Suppl. Fig 2). These observations led us to assess whether Ase promotes NB formation in a cell autonomous or non-cell autonomous manner. Thus, we produced small clones expressing high levels of Ase and we observed ectopic expression of the NB marker Mira in 58.8% of the clones originated within the NE. These ectopic Mira^+^ cells were always located within the clones but seemed to have delaminated from the NE just as NBs do (Fig. 3E,E’). Remarkably, no substantial alteration in the pattern of expression of L’sc was evident when Ase was misexpressed in the NE (Fig. 3E,E′; Suppl. Fig 2C,D). Together, these results demonstrate that Ase is sufficient to transform NE cells into NBs and they strongly suggest that this action is not mediated by L’sc.

### L’sc is not sufficient to promote NE-NB transition

It has been reported that the transient expression of the proneural factor L’sc signals the transition of NE cells to NBs in the OPC. Furthermore, it seemed that L’sc expression was sufficient to induce NBs and it was necessary for the timely onset of NB differentiation (Yasugi et al., 2008). Strikingly, we found that L’sc is still detected transiently in individual cells in *ase*^*1*^ brains (Fig. 2B) despite the strong alterations observed in the NE-NB transition (Fig. 2B,D and E). Moreover, our data indicate a very strong induction of the NE-NB transition by Ase misexpression (Fig. 3). Therefore, the capacity of L’sc to induce NB formation was compared to that of Ase by performing *l’sc* misexpression in NE cells under the same conditions as those we performed with Ase. When strong *l’sc* expression was driven in the NE for 36h, very few ectopic cells expressing NB markers were observed within the NE. Moreover, no obvious changes in the structure or size of the NE or OL were detected relative to control brains (Fig. 4A-D; Suppl. Fig. 3). Furthermore, a short (8h) induction of L’sc in the NE did not trigger NB marker expression by the NE (Figure 4E-F). Therefore, we concluded that Ase misexpression in the NE clearly has a more robust proneural effect than that of L’sc.

**Figure 4.**
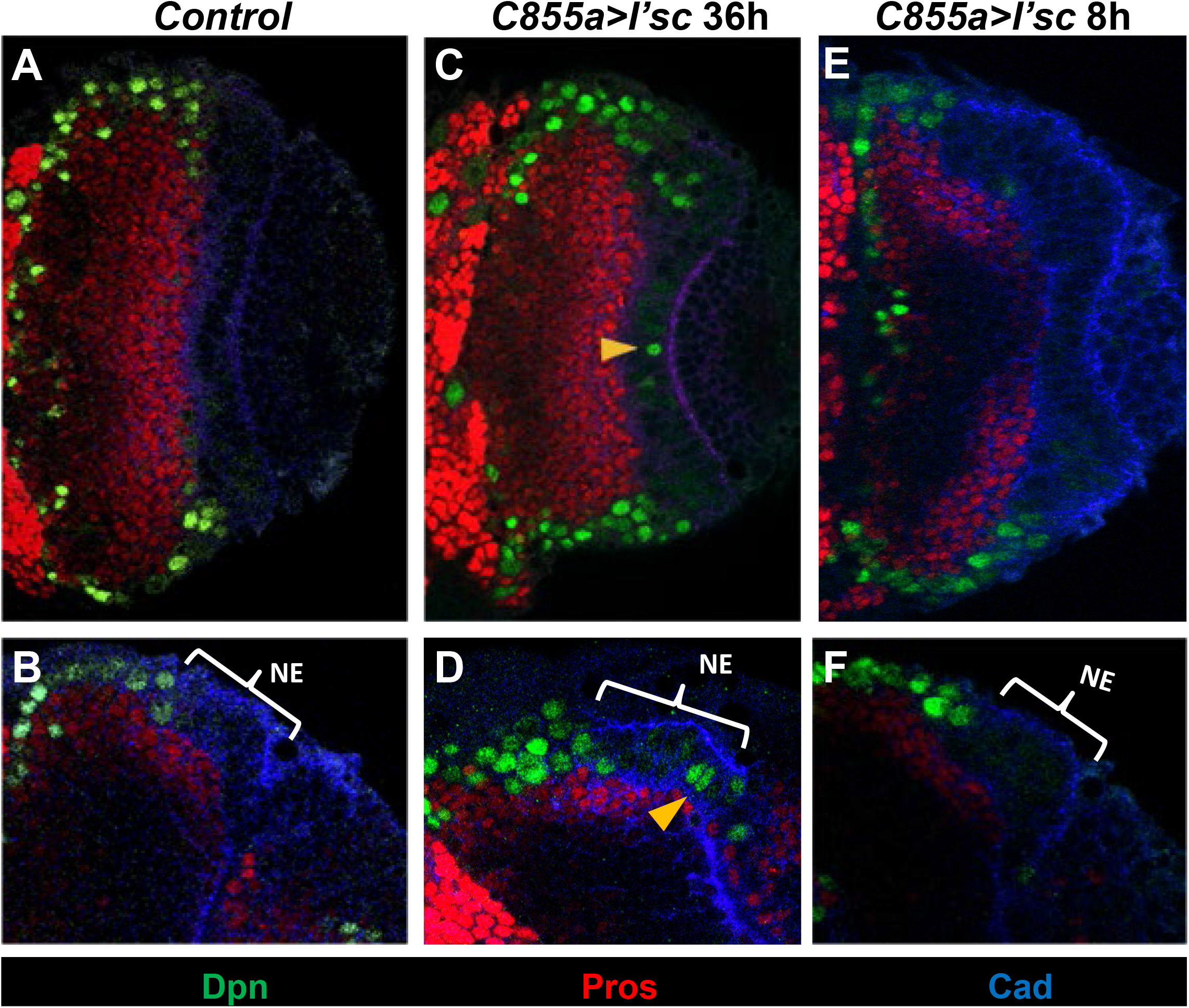
L’sc misexpression in the NE generates very few NBs. Confocal images acquired close to the surface (**A, C**) or from a deep layer (**B, D**) of the OPC of control (*c855 Gal4*) and *c855 Gal4/UAS L’sc* larval brains after a 36h induction. Note that compared with *ase* misexpression (Fig. 3 A,B) the NE has a relatively normal shape and a very few Dpn^+^ cells are located inside it (yellow arrowheads). **E, F**. There are no Dpn^+^ cells in the NE after *L’sc* misexpression (*c855 > L’sc*) for 8h compared to the large number observed after *Ase* misexpression in the same time period (Fig. 3D).

### L’sc promotes Ase expression in a non-cell autonomous manner during the NE-NB transition

Considering that the transient expression of *l’sc* precedes the peak of Ase during the NE-NB transition, we wondered if L’sc might be involved in upregulating *ase* expression. To address this issue, we analyzed *ase* expression in brains where *l’sc* was misexpressed in the NE using the *c855a-Gal4* line and we found very few scattered Ase-expressing cells within the NE (Fig. 5A-B, yellow arrowhead). By contrast, when ectopic *l′sc* expression was induced in OPC NBs using the *ase-Gal4* driver (see Suppl. Fig. 1C), Ase staining increased dramatically in all NBs, such that the peak of Ase was not evident in the transition cells (Fig. 5C). We next examined whether this activation of Ase by L’sc occurs in a cell- or non-cell autonomous manner by inducing *l′sc* expression in small clones, paying particular attention to clones located within the NE or at its medial border. We only observed an increase in Ase expression at the outer border of the clones and always outside the NE (Fig. 5D,E and E’). Furthermore, when the clone covered the transition cells, the native expression of *ase* was abolished within the clone (Fig. 5E,E’). Together, these results indicate that although L’sc alone cannot promote Ase expression in NE cells it can induce it in transition cells in a non-cell autonomous manner.

**Figure 5.**
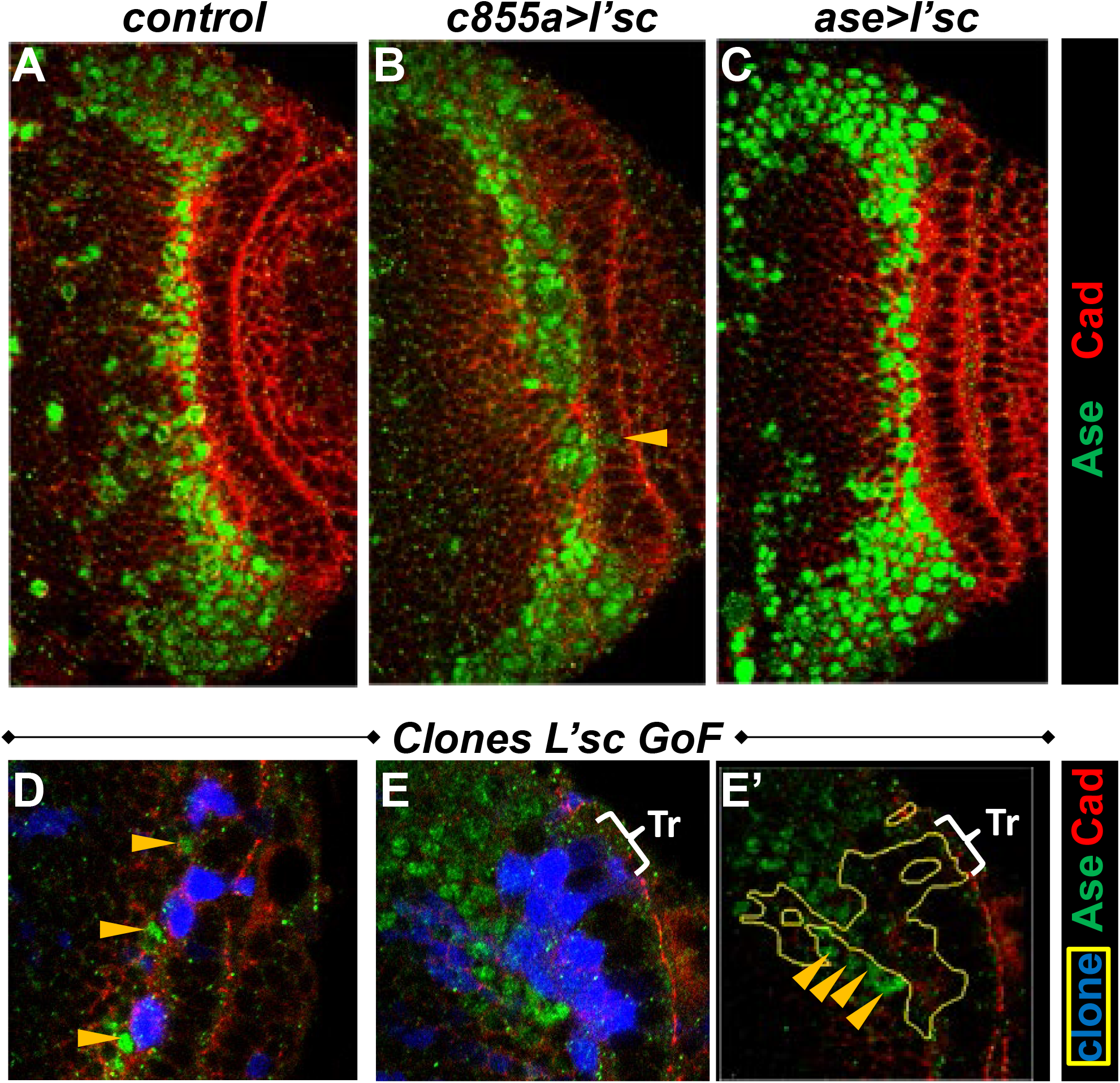
Effects of L’sc GoF on Ase expression at the NE-NB transition. **A-C**. Confocal images acquired close to the surface of the OPC of control (c855a Gal4), *c855a Gal4/UAS l’sc* and *Ase Gal4/ UAS l’sc* larval brains after a 14h induction. **B**. Note that only one Ase^+^ cell (arrowhead) is detected in the whole NE after the misexpression of *l’sc*. **C**. The intensity of Ase labeling in NBs increases strongly after the misexpression of *l’sc* relative to the control NBs (**A**). **D, E**. Clonal analysis of *l’sc* misexpression. **D**. High magnification confocal detail of the ventral surface in which three small clones are located at the edge of the NE. Note that some Ase^+^ cells (arrowheads) are situated at the outer medial border of the clones. **E**. A large clone located in the NE and in the transition region (Tr) does not contain Ase^+^ cells. Remarkably, the native *Ase* expression disappears in the Tr. There are several Ase^+^ cells (arrowheads) located outside the clone, adjacent to its inner border.

### Notch activity is necessary and sufficient for *ase* upregulation at the NE-NB transition

As mentioned previously, Notch signaling is fundamental for the progression of the proneural wave and it follows a very dynamic pattern during the NE-NB transition (Contreras et al., 2018; Egger et al., 2010; Ngo et al., 2010; Orihara-Ono et al., 2011; Reddy et al., 2010; Wang et al., 2011b; Weng et al., 2012; Yasugi et al., 2010). Notch activity is linked to *l’sc* expression, which promotes cell-autonomously the expression of *Dl* that in turn activates Notch signaling in the cells adjacent to the L’sc^+^ transition cell (Fig. 6A). Thus, we wondered if Notch signaling could regulate *ase* expression at the NE-NB transition.

**Figure 6.**
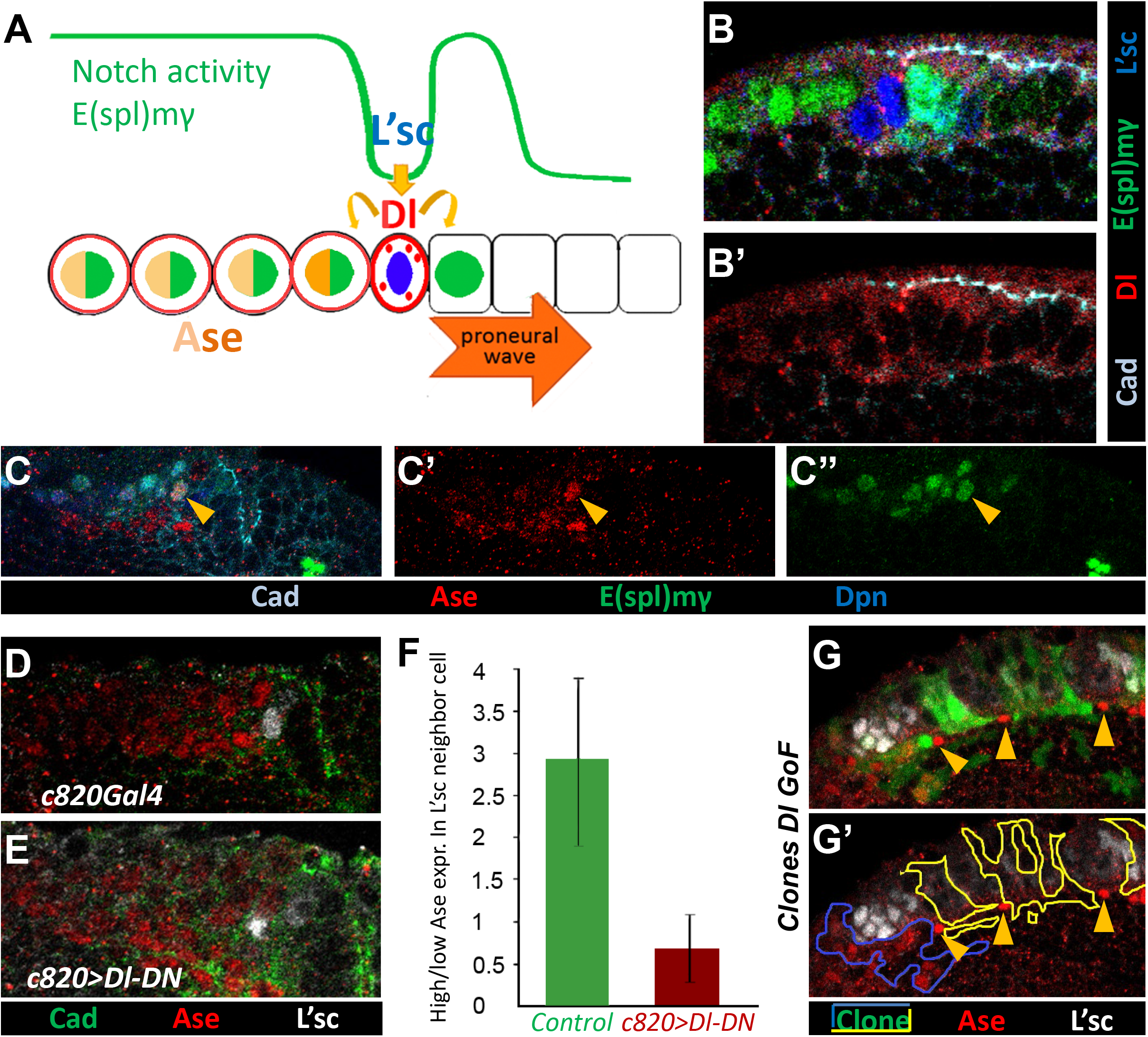
Effects of Notch GoF and LoF on Ase expression. **A.** Scheme of Notch signaling at the NE-NB transition. *l’sc* expression promotes the expression of *Dl* that activates Notch signaling (i.e. E(spl)mγ) in the neighboring cells. In the NBs, Notch activity is maintained by low *Dl* expression in all cells. **B.** Activation of Notch signaling as depicted by E(spl)mγ-GFP reporter on both sides of the strong Dl^+^/L’sc^+^ transition cells. **C, C’, C’’**. The peak of Ase cell (arrowhead) exhibit strong Notch signaling. **D, E**. The OPC of *c820 Gal 4 UAS Dl-DN* larvae show a strong decrease of Ase labeling in the L’sc^+^ neighbor cells relative to the control. **F**. Analysis of the ratio of L’sc neighboring cells with strong and weak Ase labeling in control (n=5) and *c820>Dl-DN* brains (n=12), displaying a significant difference (p-value=0.018, ***t***-student test). **G-G’**. Several Dl GoF clones located at the NE (yellow line) lack Ase immunostaining but there are strongly Ase labelled cells (yellow arrowheads) located adjacent to them. By contrast, a clone located in the NB region (blue line) contains many Ase expressing cells.

To evaluate this possibility, we first studied the correlation between *ase* expression and Notch activation in the OL using an E(spl)mγ reporter that drives GFP expression in cells in which this pathway is activated (Almeida & Bray, 2005). Thus, Notch activation was typically found on both sides of the Dl^+^/L’sc^+^ transition cells and in NBs (Fig. 6B-B’), and remarkably, we observed strong Notch activity in the peak of Ase cells (Fig. 6C). This suggested that Notch signaling could promote *ase* expression at the NE-NB transition. To test this possibility further, we blocked Notch activity by expressing a dominant negative Delta construct (Dl-DN) that produces a truncated Dl protein that interacts intracellularly with Notch, preventing it from reaching the surface of the cell and thereby blocking Notch activity in a cell autonomous manner (Huppert et al., 1997). When Dl-DN expression was induced in NE and transition cells using the *c820-Gal4* line, *Ase* expression was severely dampened in the cells neighboring the L’sc^+^ transition cell (Fig. 6D-F). Hence, the downregulation of Notch activity suppressed the peak of Ase at the NE-NB transition. Conversely, when we induced ectopic Notch activity in *Dl GoF* clones located in the NE, ectopic Ase-expressing cells were only observed at the outer border of these clones, at the edge of the NE (Fig. 6G,G’). Hence, the activation of Notch signaling by Dl at the NE border appears to induce ectopic peaks of Ase expression, something reminiscent of the effect of L’sc-expressing clones (Fig. 5D,E). By contrast, Dl expressing clones located in the NB region contained *ase* expressing cells (Fig. 6G’). Together, these findings indicate that Notch activity is necessary and sufficient to induce *ase* expression at the NE-NB transition.

Since Dpn seems to be upregulated in OPC NBs immediately after the peak of Ase, we wondered if Dpn could be involved in the downregulation of Ase in NBs after its peak in the transition cells. To this end, we performed RNAi of *Dpn* using *the c820-Gal4* line and while this extensively depleted Dpn in NBs, it did not apparently enhance *ase* expression (Suppl. Fig.4). Hence, Dpn does not seem to be required to repress *ase* in NBs.

## DISCUSSION

### Ase is necessary and sufficient to promote the NE to NB transition

Among the key players regulating events in neurogenesis, the proneural bHLH TFs deserve special attention as they fulfil major, evolutionary conserved roles. Studies in *Dm* and vertebrates have shown that these factors are necessary and sufficient to initiate a developmental program that generates NPs committed to produce neuronal and glial lineages (reviewed by Bertrand et al., 2002; Campos-Ortega, 1993; García-Bellido & de Celis, 2009; Guillemot & Hassan, 2017; Jan & Jan, 1993; Ross et al., 2003; Wilkinson et al., 2013). In *Dm,* the proneural members of the AS-C, *ac, l’sc* and *sc* play a key role in driving epidermal cells towards a neural fate. Thus, they are expressed in clusters of ectodermal cells (proneural clusters) and they promote the formation of sensory organ precursors (SOPs) in the embryonic and adult PNS, and of NBs in the embryonic CNS (reviewed by Campos-Ortega, 1993; García-Bellido & de Celis, 2009; Jan & Jan, 1993). Initial studies indicated that *ase*, the fourth member of the AS-C, was functionally different since it is not expressed in the ectoderm but rather in SOPs and their lineages, where it persists longer than the proneural AS-C factors (Alonso & Cabrera, 1988; Brand et al., 1993; González et al., 1989; Jarman et al., 1993). Moreover, *ase* deletion caused abnormal differentiation of sensory organs (Domínguez & Campuzano, 1993). Accordingly, *ase* seemed to be a neural precursor gene rather than a proneural gene. Nevertheless, Ase seems to play a proneural role in the generation of some wing margin bristles (Domínguez & Campuzano, 1993) and it also displays proneural potential since its ectopic expression is capable of initiating sense organ fate (Brand et al., 1993; Domínguez & Campuzano, 1993).

In the developing larval CNS, *ase* was found to be expressed in NBs and their progeny (Brand et al., 1993; González et al., 1989; Yasugi et al., 2008), which in principle also pointed to a neural precursor gene role. Indeed, Ase has been used extensively as a Type I NB marker. Nevertheless, we previously found that *ase* was transiently upregulated in GCs generated by Type I and OPC NBs, playing a key role in the mechanisms that induce the cell cycle exit and terminal differentiation of these neuronal precursors (Shaikh et al., 2016). Interestingly, this function very much resembles the classic role played by the AS-C vertebrate orthologues Ascl1/Mash1 and Neurogenins in promoting the cell cycle arrest and neuronal differentiation of NPs (reviewed by Bertrand et al., 2002; Castro et al., 2011; Wilkinson et al., 2013).

We have shown here that *ase* is upregulated at the NE-NB transition of the OPC. Two types of intermediate NPs have been described previously as this NE-NB transition progresses: PI progenitors are defined by the expression of NE markers and strong Notch activity, whereas PII progenitors, while still exhibiting NE markers, are distinguished by strong *l’sc* expression and weak (or no) Notch activity (Sato et al., 2013; Yasugi et al., 2010). Thus, the peak of Ase appears to identify a third step in this transition (PIII) after the transient expression of *l’sc*, since these cells strongly expressing *ase* have lost NE makers, they receive strong Notch signal and barely exhibit NB markers (Fig. 7A,B).

**Figure 7.**
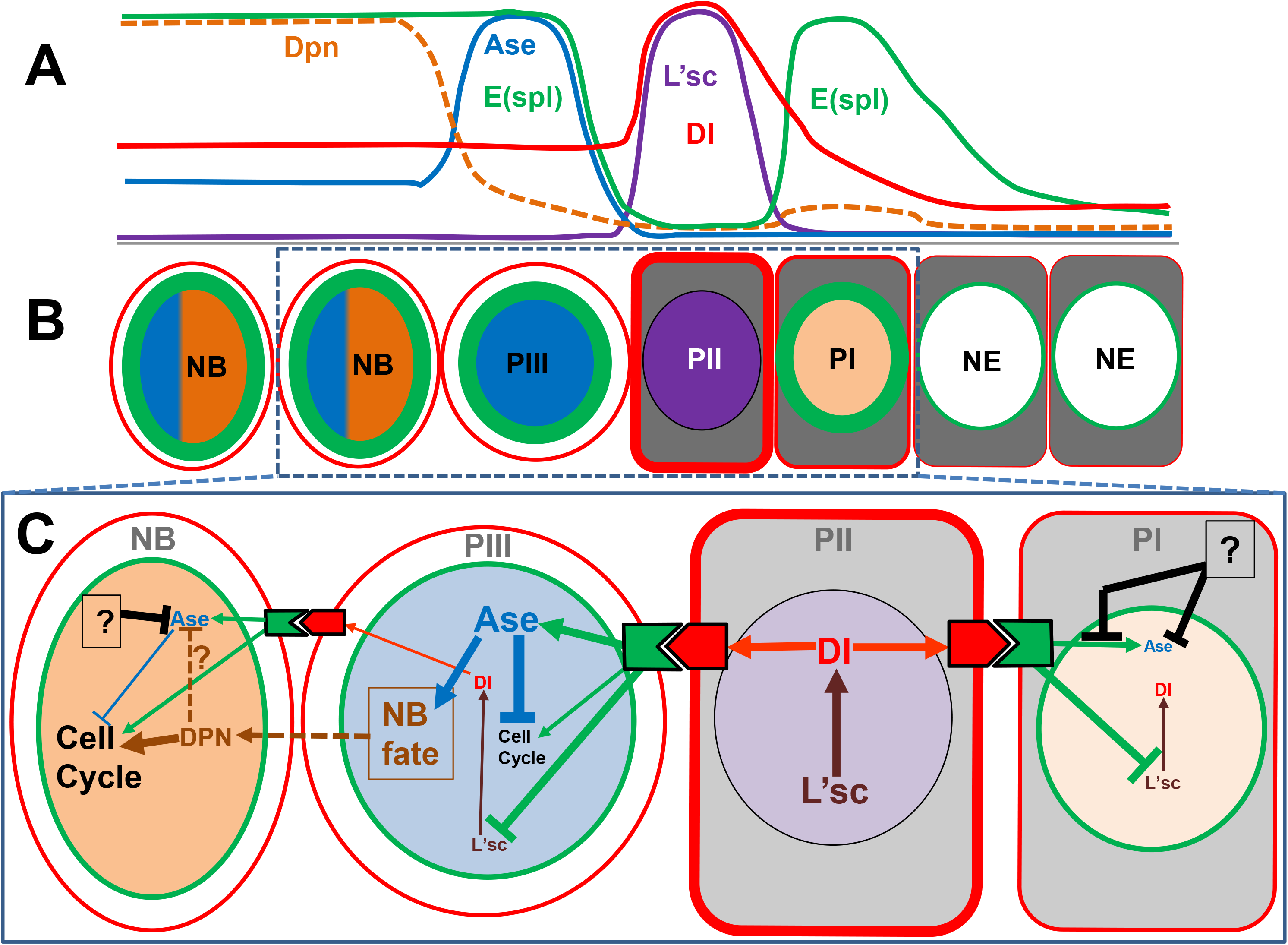
Working model: integration of Ase proneural function with L’sc and Notch signaling promotes the NE-NB transition. **A**: Schematic representation of the expression profiles of Ase, Dl, Dpn, L’sc and E(spl)mγ (Notch signal activity) during the NE-NB transition. **B**. Distinct cellular stages along the NE-NB transition. The nuclear and cell wall colors correspond to Ase, Dl, Dpn, L’sc and E(spl)mγ as in panel A. **C**. Working model with the interactions of Ase, Dl, Dpn, L’sc and Notch activity facilitating the NE-NB transition. The “?” elements refer to unknown mechanisms of *ase* repression.

Most importantly, our present results demonstrate that this upregulation of *ase* is necessary and sufficient to promote a correct NE to NB transition, providing the first direct evidence of a proneural role for Ase in CNS neurogenesis. Notably, our data shows that the LoF of *ase* does not fully impede the NE-NB transition. Thus, in conjunction with the existing data generated using AS-C deficiencies (Yasugi et al., 2008), our results strongly suggest that this key transition is very robust and possibly relies on redundant mechanisms.

In summary, the present results together with our previous data (Shaikh et al., 2016; Shaikh & Tejedor, 2018) demonstrate that Ase plays major roles in regulating the program of neurogenesis in the Drosophila OL by promoting the differentiation of NPs through successive phases: First, from proliferative (NE) to self-renewing (NBs) NPs and then, to the cell cycle exit and terminal differentiation of neuronal precursors (GCs).

### Similar and divergent functions of Ase and its vertebrate orthologue Ascl1/Mash1

Remarkably, our results are also consistent with the classic actions of *Ascl1* at sequential stages of vertebrate CNS neurogenesis, specifying NP subtypes in addition to cell cycle exit and terminal differentiation of neurons (reviewed by Bertrand et al., 2002; Castro & Guillemot, 2011; Guillemot, 2007; Guillemot & Hassan, 2017; Wilkinson et al., 2013). Nevertheless, we do not find indications that *ase* fulfils a similar role to *Ascl1* in promoting NP proliferation by activating positive cell cycle regulators (Castro et al., 2011). Indeed, while the *Ascl1* knock-out induces premature cell-cycle exit of ventral telencephalon NPs (Castro et al., 2011; Pilz et al., 2013), we have not found such an effect in null *ase* mutant or after RNAi knockdown. As a matter of fact, the peak of Ase in the NE-NB transition possibly coincides with a prolonged cell cycle of transition cells (Orihara-Ono et al., 2011; Reddy et al., 2010), which in principle points to an anti-proliferative action of Ase. It is particularly noteworthy that Ase can induce cell cycle arrest of GCs by promoting the expression of *dacapo* (*dap*) (Shaikh et al., 2016; Wallace et al., 2000), the major cyclin dependent kinase inhibitor in *Dm* (de Nooij et al., 1996; Lane et al., 1996). In that sense, the downregulation of *ase* after its expression peak could allow NBs to continue cycling. Thus, it would be very interesting to analyze in more depth the possible role played by Ase in the regulation of the cell cycle during the NE-NB transition.

It is well established that *Ascl1* expression oscillates in NPs of the mouse embryonic brain and this is crucial for its sequential roles in neurogenesis (reviewed by Imayoshi & Kageyama, 2014; Vasconcelos & Castro, 2014). Similarly, our current data also indicates that the peak (oscillation) of *ase* expression is required for the correct NE-NB transition, since flattening the peak (i.e. dampening expression in the transition cells but maintaining it in NBs) by RNAi knockdown produced the same alterations as the *ase* deletion. Nevertheless, there are several reasons to think that this wave of Ase in the OPC differs from the oscillatory expression of *Ascl1* in the embryonic ventral telencephalon. In this tissue, this oscillation of *Ascl1* expression is associated with its role in maintaining a proliferative/self-renewal state, while its sustained expression correlates better to terminal differentiation (reviewed by Imayoshi & Kageyama, 2014; Vasconcelos & Castro, 2014). By contrast, we have no evidence that the sustained expression of *Ase* in either NEs (this work) or NBs (Shaikh et al., 2016) leads to neuronal differentiation. Conversely, it appears that the transient expression of *ase* during the NE-NB transition (herein) and in newborn GCs (Shaikh et al., 2016) is what drives NB specification and neuronal differentiation, respectively.

An additional reason to think that the oscillations of Ase and Ascl1 are functionally different is that Dpn does not appear to be fully required for the downregulation of *ase* after its expression peak. This is in contrast to the well-known repression that the Dpn orthologue Hes1 exerts on *Ascl1* expression during its oscillation in the ventral telencephalon. This repression is required to maintain NPs in a self-renewing state (reviewed by Imayoshi & Kageyama, 2014). Nevertheless, there is evidence that Dpn can repress *ase*, as Dpn consensus sites have been found in the *ase* gene (Southall & Brand, 2009). More importantly, we previously found that the overexpression of Dpn substantially reduced *ase* expression in NBs (Shaikh & Tejedor, 2018). Nevertheless, we cannot rule out that the lack of effect of Dpn knockdown on *ase* expression could be due to functional redundancy with other bHLH genes that might substitute for it, as reported in other cell contexts (Zacharioudaki et al., 2012).

In summary, Ase and Ascl1 seem to fulfil equivalent and divergent functions in brain neurogenesis. We believe that the main motive underlying these differences resides in the divergent strategies adopted by genetic programs to the different cell patterns of neurogenesis: salt and pepper (mouse telencephalon) vs. expression wave across the tissue (*Dm* OPC).

### Ase cooperates with Notch signaling and L’sc to ensure a timely NE-NB transition

Elucidating the genetic programs and molecular mechanisms that control neurogenesis has been a major focus of research over the past three decades (Doe, 2008; Götz & Huttner, 2005; Guillemot, 2007b; Hartenstein & Stollewerk, 2015; Homem et al., 2015; Taverna et al., 2014). Consequently, many genes and individual mechanisms have been identified that operate at different stages. Nevertheless, how these mechanisms are integrated to coordinate the whole neurogenic process remains poorly defined.

The NE-NB transition is a good example of a process regulated by multiple signaling pathways, although it is only just beginning to be understood how they are coordinated. For instance, Notch signaling seems to be involved in several sequential steps in this transition. Initially, it is fundamental to maintain NE cell fate and proliferation (Egger et al., 2010; Ngo et al., 2010; Orihara-Ono et al., 2011; Reddy et al., 2010; Wang, Liu, et al., 2011; Yasugi et al., 2010). Similarly, the Notch pathway plays a key role in maintaining NPs undifferentiated during the early neurogenic phases of the developing vertebrate CNS (reviewed by Artavanis-Tsakonas et al., 1999; Imayoshi et al., 2010; Mizutani & Saito, 2005). Nevertheless, it is well known that the effects of Notch signaling depend on the cell context (reviewed by Bray, 2006; Kageyama et al., 2009; Koch et al., 2013; Louvi & Artavanis-Tsakonas, 2006; Pierfelice et al., 2011). Thus, it has become clear that Notch signaling is also involved in promoting the transformation of NE progenitors into neurogenic NPs (Hämmerle & Tejedor, 2007; Hatakeyama & Kageyama, 2006; Kageyama et al., 2009). This was evident in the prospective spinal cord of the chick embryo where a caudo-rostral wave of Delta-Notch signaling in proliferating NPs mediates their transition to neurogenesis (Hämmerle & Tejedor, 2007). This action resembles what happens in the proneural wave at the *Dm* OL in which Notch activity is essential for its progression, as discussed above. Indeed, Notch LoF causes premature transformation of NEs cells into NBs (Egger et al., 2010; Ngo et al., 2010; Orihara-Ono et al., 2011; Reddy et al., 2010; Wang, et al., 2011b; Yasugi et al., 2010). Conversely, ectopic activation of Notch signaling delays the NE-NB transition (Wang, Liu, et al., 2011; Yasugi et al., 2010). Finally, at the end of the transition Notch signaling is also essential for NB proliferation (Zacharioudaki et al., 2012; Zhu et al., 2012).

However, it is less clear how Notch signaling acts at the intermediate steps between the NE and NB. As described above, Notch activity is very dynamic during the NE-NB transition and thus, it is weakly activated across most of the NE while its activity is very strong in the most medial NE cells (PI transition progenitors) that are neighbors of the high Dl^+^/L’sc^+^ PII transition cells in which the Notch pathway is inactive. Finally, Notch signaling is again upregulated in the adjacent PIII transition cell and the NBs (Fig. 7A). The results we present here demonstrate that Delta-Notch signaling is necessary and sufficient to promote *ase* expression in the PIII progenitors. This reveals a novel context specific role for the Notch pathway in neurogenesis since in contrast to the positive effect on *ase* expression seen here, Notch signaling represses proneural gene expression in classic neurogenic models by different means and NP production is in general associated to low Notch activity (Artavanis-Tsakonas et al., 1999; Bertrand et al., 2002). Nevertheless, our Dl and L’sc GoF results indicate that *ase* is repressed strongly in the NE by an unknown mechanism that serves to prevent its premature upregulation. Thus, despite the strong Notch signals available to the most medial NE cells (i.e. PI progenitors), *ase* is only activated by Notch at the most medial transition cells (i.e. PIIIs) after *l’sc* is expressed in PIIs (Fig. 7A-C).

Although the *ase* gene promoter contains binding sites for AS-C/daughterless heterodimers (Jarman et al., 1993) and *ase* appears to lie downstream of proneural genes in PNS neurogenesis (Brand et al., 1993), our data indicate that the action of L’sc on *ase* expression in the OL is not cell autonomous but rather, it is mediated by the activation of Notch signaling in the adjacent cell. This could be one of its main functions in this context given that L’sc, in contrast to Ase, does not appear to be sufficient to drive the NE-NB transition. In addition, L’sc is also involved in promoting the transition from the PI to PII progenitor state by inducing the expression of Neuralized (Contreras et al., 2018). Through these synchronized functions, L’sc could control the timing of NE-NB progression.

In addition to the cell autonomous promotion of neurogenesis, neuronal commitment, cell cycle exit and terminal differentiation, it is well stablished that the expression of proneural factors by NPs activates Notch signaling in the adjacent progenitors (reviewed by Bertrand et al., 2002; García-Bellido & de Celis, 2009; Kageyama et al., 2009). Our results support the idea that these two parallel proneural functions have been split between L’sc and Ase in the case of the NE-NB transition. Thus, L’sc appears to be in charge of activating Notch in the adjacent transitional NP (PIII), while Ase promotes the differentiation of these cells into NBs (the neurogenic progenitors).

In summary, our results reveal a novel and crucial proneural function of Ase in CNS neurogenesis, which is integrated with Notch signaling and L’sc actions promoting the progressive transformation from NE cells into NBs.

## MATERIALS AND METHODS

### Drosophila melanogaster strains

Fly stocks were raised at 25 ºC on standard medium. The following strains were used: *Oregon R* (wt); *ase*^*1*^ (formerly known as *sc*^*2*^, González et al., 1989); *c855a-Gal4* (Hrdlicka et al., 2002); *c820-Gal4* (Manseau et al., 1997); *ase-Gal4* (San-Juán & Baonza, 2011); *UAS-ase-RNAi* (VDRC stock # GD12444; Dietzl et al., 2007); *UAS-ase* (Brand & Dormand, 1995); *UAS-L’sc* (Carmena et al., 1995); *UAS-dGFP* (Lieber et al., 2011); *hs FLP; act-FRT-y+-FRT Gal4 UAS GFP* (BDSC stock # 30558); *UAS Dl-DN*(BDSC stock # 5613, Huppert et al., 1997), *UAS-Dl* (BDSC stock # 26694) and *UAS-dpn-RNAi* (BDSC stock # 26320; Perkins et al., 2015).

### Gene misexpression and RNAi using the UAS/Gal4 system

Larvae carrying Gal4/UAS constructs were kept at a restrictive temperature (17 ºC) until the appropriate stage of development and induced at a permissive temperature (29 ºC) until the wandering stage, at which they were dissected. RNAi induction was maintained for 24h, while *ase* and *l’sc* misexpression was induced over 8h at the late 3^rd^ instar, or for 36h at around the beginning of 3^rd^ instar, as indicated in each experiment.

### Generation of clones

Recombinant clones were generated using the Flip-out technique (Struhl & Basler, 1993). To this end, *hsp70-Flp; Actin5C <yellow, [stop]> Gal4, UAS-GFP* female flies were crossed with males carrying the different UAS constructs indicated in each experiment. Larvae were raised at 25 ºC and clones were generated by inducing FRT-mediated recombination with a 15 min heat shock at 37 ºC. Subsequently, the larvae were incubated at 29 ºC for 16h before dissection.

### Immunohistochemistry

Third instar larval brains were dissected out in PBS and fixed in 4% paraformaldehyde (PFA) for 20 min at room temperature (RT), or for 1h at 4 ºC (for Dl immunostaining). Brains were then washed with PBST-0.5% (0.5% Triton X-100 in PBS) and non-specific binding was blocked with 10% normal goat serum (NGS) in PBST-0.5% + 0.02% sodium azide for 1h (or 4h for Dl immunostaining) at RT. The brains were probed overnight with the primary antibodies at 4 ºC and for 1h at RT the next day. After washing in 0.5% bovine serum albumin (BSA) in PBST-0.2%, the brains were then incubated for 1-2h at RT with fluorescent conjugated secondary antibodies (Alexa-488, Alexa-594, Alexa-647, Cy3 or Cy5: Jackson ImmunoResearch) diluted 1:500. Samples were then washed sequentially at RT with BSA/PBST-0.2%, PBST-0.3%, and PBS for 30 min each, and finally mounted in Vectashield H-100 medium (Vector Lab, Germany). Immunostaining was examined by confocal microscopy (Olympus FV10i fluoview or Zeiss LSM 880-Airyscan Elyra PS.1.), analyzing the images with FV10-ASW 4.2 viewer or ZEN lite 3.2 blue edition.

The following antibodies were used: rat anti-Cad (1:120, DSHB, Clone DCAD2: Oda et al., 1994); mouse anti-Mira (1:100, a generous gift of F. Matsuzaki, Clone PLF8); mouse anti-Pros (1:30; DSHB clone MR1A: Campbell et al., 1994); guinea pig anti-Dpn (1:3000, a generous gift of A Carmena: Rives-Quinto et al., 2017); rabbit anti-Ase (1:2000, a generous gift of Y.N. Jan: Brand et al. 1993); guinea pig anti-L’sc (1:1500, a generous gift of M. Sato: Suzuki et al., 2013); mouse anti-Dl (1:30, DSHB, Clone C594.9B: Sun & Artavanis-Tsakonas, 1996); chick anti-GFP (1:3000: AvesLab).

### Quantification and statistics

The number of labelled cells was quantified manually using the FV10-ASW 4.2 viewer software and analyzed statistically with Microsoft Excel 2010. The data were evaluated with ***t*** student tests for two-tailed heteroscedastic distributions. Differences were considered significant at p < 0.05. Mira and Cad co-expressing cells were counted at the medulla NE to NB transition in serial confocal sections every 2 μm in the 30-60 μm Z-axis interval from the ventral surface of the OL.

To assess the changes in expression, control and mutant samples were processed in parallel using the same solutions. Moreover, all the samples in each experiment were analyzed during the same confocal microscope work session, using the same optical and image acquisition parameters.

## ABBREVIATIONS

*I*: current (A)
AS-C: achaete scute complex
bHLH: basic helix-loop-helix
BSA: bovine serum albumin
CB: central brain
CNS: central nervous system
Dm: Drosophila melanogaster
GC: ganglion cell
GMC: ganglion mother cell
GoF: gain of function
IHC: immunohistochemical
IPC: inner proliferation center
LoF: loss of function
NB: neuroblast
NE: neuroepithelial/neuroepithelium
NGS: normal goat serum
NP: neural progenitor
OL: optic lobe
OPC: outer proliferation center
PBS: phosphate buffered saline
PFA: para-formaldehyde
PNS: peripheral nervous system
RT: room temperature
SOPs: sensory organ precursors
SVZ: subventricular zone
TF: transcription factor
VZ: ventricular zone

## ACKNOWLEDGEMENTS

We are very grateful to S. Campuzano for useful suggestions and critically reading of the manuscript. We are indebted to A. Baonza, H. Bellen, S. Campuzano, A. Carmena, C. Doe, M. Dominguez, C. Estella, C. Gonzalez, Y-N. Jan, A. Jarman, F. Matsuzaki, M. Sato, S. Selleck, J. Skeath, G. Struhl, H. Vaessin, the Vienna Drosophila Resource Centre (VDRC), the Bloomington Stock Center (BSC), and the Developmental Studies Hybridoma Bank (DSHB) for providing us with flies, antisera and other molecular tools.

## AUTHOR CONTRIBUTIONS

M.N.S., and F.G.A. performed experiments and analyzed data.

M.M. performed experiments, analyzed data and wrote the manuscript.

F.J.T. developed the concepts, performed experiments, analyzed the data and wrote the manuscript.

## DATA AVAILABILITY

Datasets supporting this work will be deposited in DIGITAL.CSIC (http://digital.csic.es/?locale=en) after acceptance of the paper and made publicly available at the time of publication.

---

**Suppl. Figure 1.**
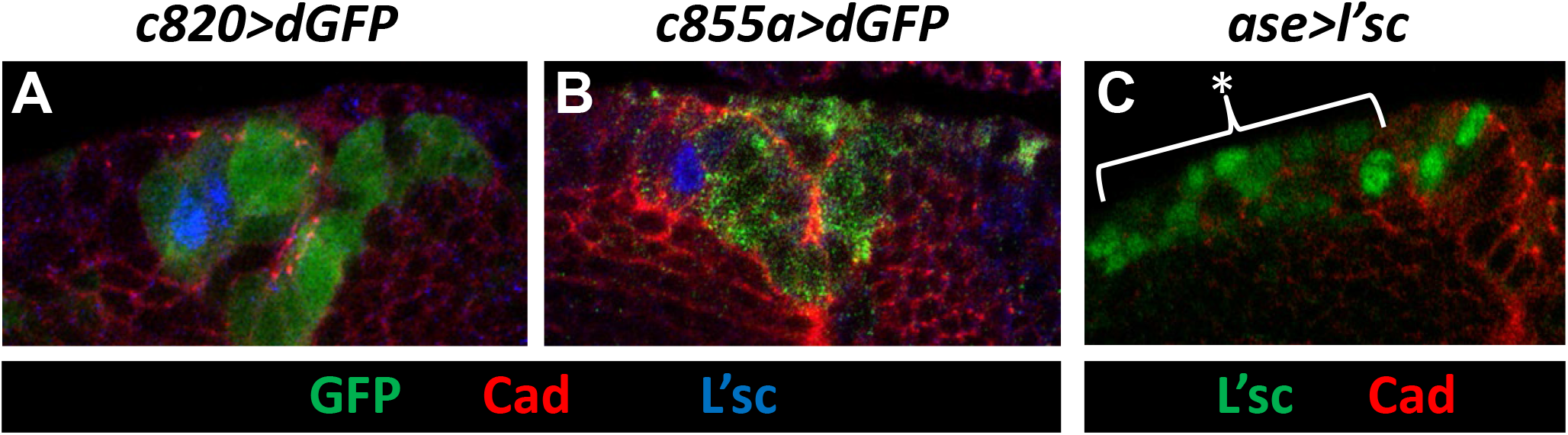
Expression patterns of various Gal4 drivers at the NE-NB transition. **A, B.** GFP expression in the OPC (deep layer) of *c820 Gal4/UAS dGFP* and *c855a Gal4/UAS dGFP* larval brains. In the *c820>GFP* sample, GFP is present in the transition cell (L’sc^+^) and in the most medial NE, while the *c855a>GFP* specimen exhibits GFP in all NE cells but not in transition cells. **C.** Expression of L’sc in the OPC (deep layer) of an *Ase Gal4/UAS L’sc* larval brain. Note the ectopic expression all along the NB region (*).

**Suppl. Figure 2.**
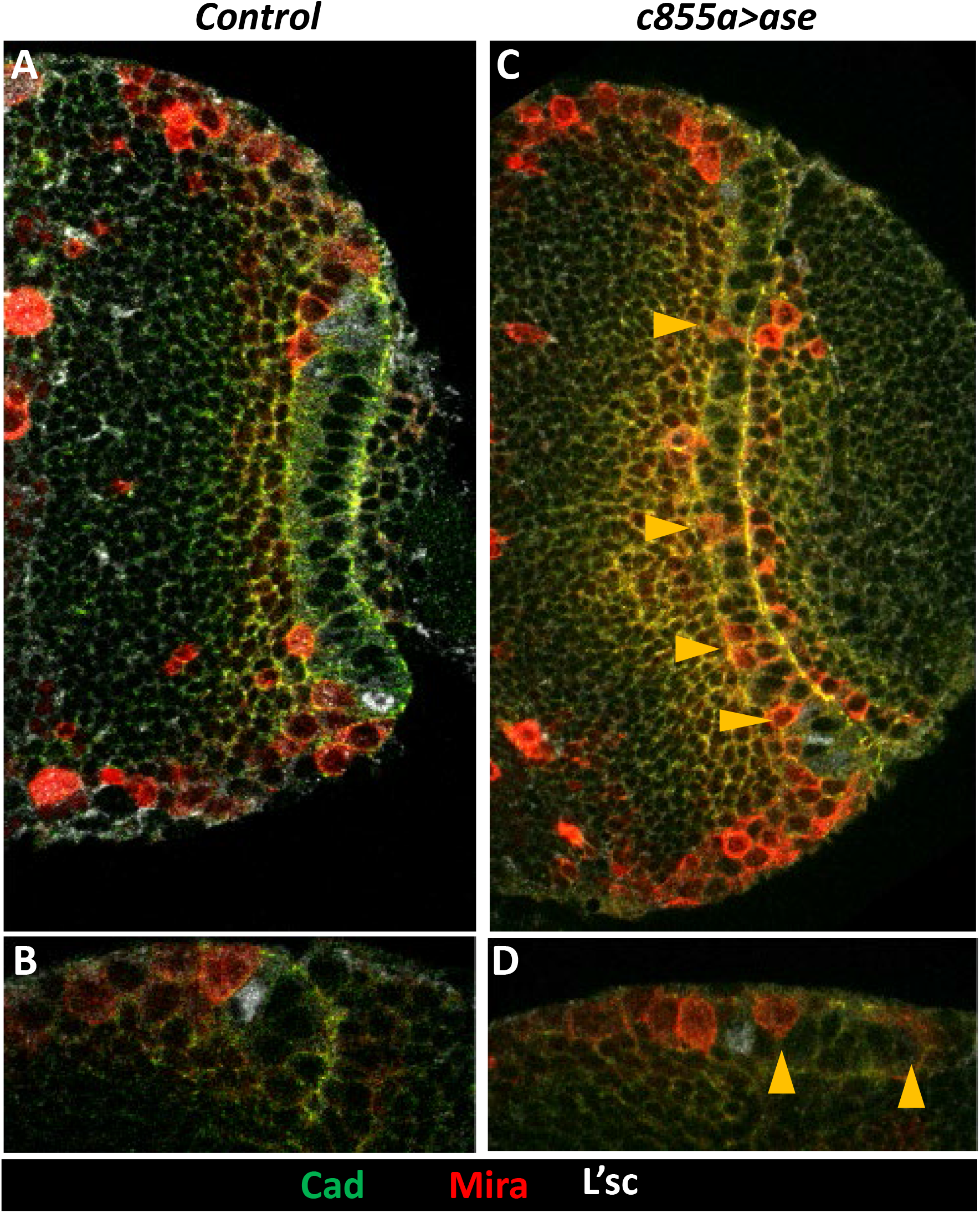
The GoF of Ase in the NE promotes the expression of Mira. Confocal images taken close to the surface (**A,B**) or in deep layers (**C,D**) of the OPC of control (*c855a Gal4*) and *c855a Gal4/UAS ase* larval brains after a 8h induction. Note the presence of Mira^+^ cells intermingled in the NE (yellow arrowheads) of the *c855>ase* specimen compared to the control.

**Suppl. Figure 3.**
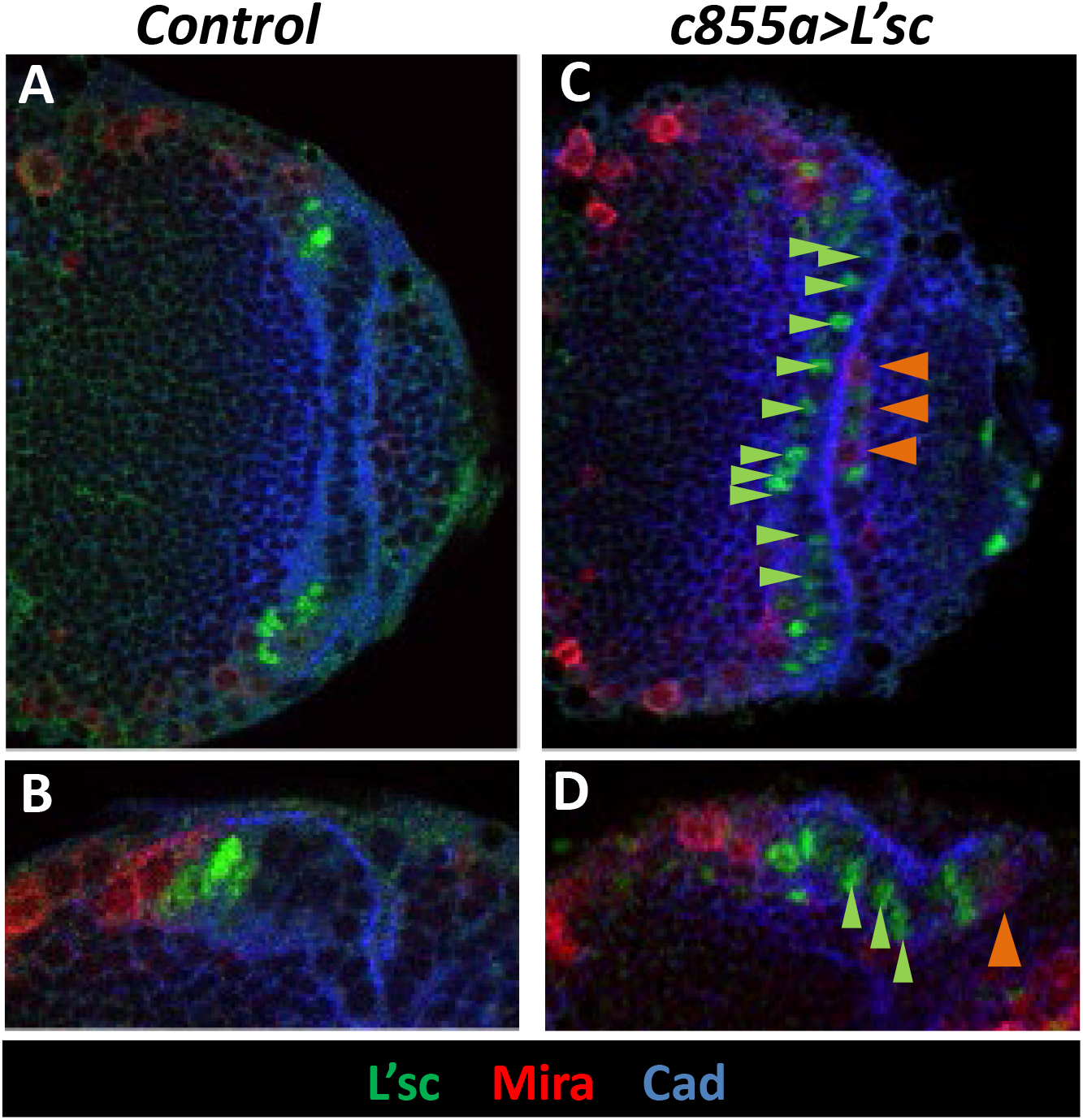
The GoF of L’sc in the NE does not promote Mira expression. Confocal images taken close to the surface (**A,B**) or in deep layers (**C,D**) of the OPC of control (*c855a Gal4*) and *c855a Gal4/UAS l’sc* larval brains after a 16h induction. Note that despite the large number of L’sc^+^ cells (green arrowheads) inside the Medulla NE (left side) of the *c855>ase* specimen relative to the control there are no Mira+ cells inside, although there are some in the Lamina NE (right side, orange arrowheads).

**Suppl. Figure 4.**
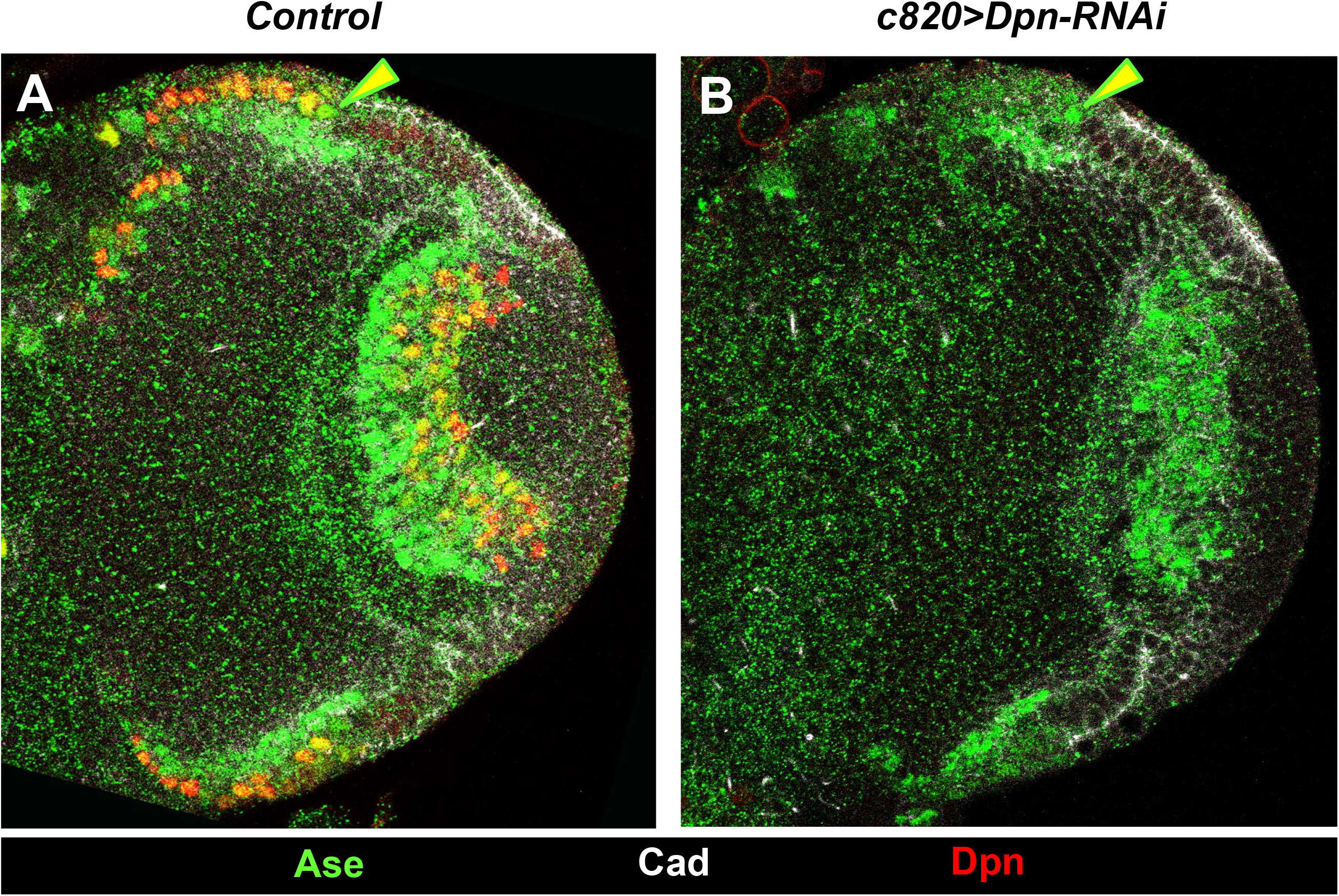
The RNAi of Dpn in NBs does not modify Ase expression. Confocal images taken at deep layers of the OPC of control (*c820 Gal4*) and *c820 Gal4/UAS Dpn-RNAi* larval brains after a 24h induction. Note that despite a severe decrease in Dpn immunostaining there is no increase in Ase labeling in the NBs relative to the control and to the Ase peak cell (arrowhead).

